# Tracing Diurnal Differences in Brain Anatomy with Voxel-Based Morphometry – Associations with Sleep Characteristics

**DOI:** 10.1101/2022.03.06.483163

**Authors:** Michal Rafal Zareba, Magdalena Fafrowicz, Tadeusz Marek, Halszka Oginska, Ewa Beldzik, Aleksandra Domagalik

## Abstract

Multiple aspects of brain functioning, including arousal, motivation, and cognitive performance, are governed by circadian rhythmicity. Although the recent rise in the use of magnetic resonance imaging (MRI) has enabled investigations into the macroscopic correlates of the diurnal brain processes, neuroanatomical studies are scarce. The current work investigated how time-of-day (TOD) impacts white (WM) and grey matter (GM) volumes using voxel-based morphometry (VBM) in a large dataset (N = 72) divided into two equal, comparable subsamples to assess the replicability of effects. Furthermore, we aimed to assess how the magnitude of these diurnal differences was related to actigraphy-derived indices of sleep health. The results extend the current knowledge by reporting that TOD is predominantly associated with regional WM volume decreases. Additionally, alongside corroborating previously observed volumetric GM decreases, we provide the first evidence for positive TOD effects. Higher replicability was observed for WM, with the only two replicated GM clusters being volumetric increases in the amygdala and hippocampus, and decreases in the retrosplenial cortex, with the latter more pronounced in individuals with shorter sleep times. These findings implicate the existence of region-specific mechanisms behind GM effects, which might be related to cognitive processes taking place during wakefulness and homeostatic sleep pressure.

## 1. Introduction

Circadian rhythmicity governs many aspects of our everyday lives. This is especially evident for processes embedded in the brain. The levels of arousal, motivation and our cognitive performance, amongst others, are all subjected to the within-day fluctuations (Marek et al. 2010; Murray et al. 2009; Saper et al. 2005). The development of chronobiology and the observation of diurnal variations at all levels of the organisation of biological systems has made us aware of an important point: physiological processes in our bodies continuously change and as such, so does the microstructural anatomy.

Investigations probing these daily fluctuations in the function and structure of the human brain have largely been made possible thanks to the use of magnetic resonance imaging (MRI). So far, the circadian effects have mostly been studied in relation to the various aspects of brain activity (Fafrowicz et al. 2019; Farahani et al. 2021; Gaggioni et al. 2014; Marek et al. 2010; Orban et al. 2020). As far as daily fluctuations in the brain structure are concerned, a few studies have reported TOD differences in cortical thickness (Elvsåshagen et al. 2017; Trefler et al. 2016) and diffusion of water molecules within white matter (WM; Jiang et al. 2014; Thomas et al. 2018; Voldsbekk et al. 2020).

To the best of our knowledge, only one work to date has investigated the TOD variability using voxel-based morphometry (VBM; Trefler et al. 2016). The technique is capable of measuring the relative volume of grey matter (GM) and WM in each voxel in the brain. Nevertheless, the discussed work restricted its analysis only to GM, reporting that in a sample of 19 individuals 4 h of controlled rest were associated with GM volume decreases in several areas within the frontal, temporal and parietal lobes, as well as in the cerebellum and caudate nucleus. Thus, currently it is unknown whether the VBM-measured WM volume is similarly affected by the TOD effect. PubMed search indicates 1734 manuscripts containing the terms “voxel-based morphometry” and “white matter”. Taking into account the popularity of the technique, it is crucial to answer the presented question. If the WM volume is indeed subjected to daily fluctuations and this factor is not properly controlled, this could contribute to erroneous findings and misleading conclusions. Furthermore, several studies have reported that variability in other circadian-related factors, such as sleep-wake homeostasis, modulates the TOD effects on brain activity (Song et al. 2019; Vandewalle et al. 2011). At present, however, it is unknown whether a similar phenomenon is present in the context of TOD differences in the brain structure.

One of the goals of the current study was to explore for the first time whether VBM-measured WM volume is subjected to daily fluctuations similar to GM. Additionally, we aimed to replicate the GM findings reported earlier in the literature (Trefler et al. 2016). The aforementioned study investigated the VBM GM TOD effect in a tightly-controlled laboratory setting. By doubling the between-session difference in circadian time to 9 h and allowing the participants to engage in their daily non-strenuous activities, we wanted to create a scenario which to a larger extent resembles the typical studies where TOD effect is not controlled at the stage of experiment planning. We hypothesised that both TOD increases and decreases in GM and WM volume could be observed as a result of, respectively, experience-dependent plasticity (Asan et al. 2021; Gibson et al. 2014; Keifer et al. 2015) and permeation of CSF-like water into the brain parenchyma (Thomas et al. 2018; Xie et al. 2013).

Furthermore, as described earlier, despite the growing interest in the subject, the relationship between sleep and TOD variability in brain structure remains uncharted. To explore this novel field, we correlated the TOD structural differences with two indices of sleep health, i.e. sleep length and sleep fragmentation. We expected the volumetric TOD increases, predominantly linked to the synaptic plasticity, to be positively associated with the sleep metrics (Bellesi et al. 2013; Li et al. 2017). As for the volumetric TOD decreases, mainly related to the diffusivity of CSF-like water, both positive and negative correlations were expected, similarly to the previously reported sleep quality-related results (Demiral et al. 2019). The current work was performed in an extended cohort (N = 72) from earlier works showing widespread TOD variation in resting-state functional connectivity (Fafrowicz et al. 2019; Farahani et al. 2021). To the best of our knowledge, it is the largest dataset used to investigate the interindividual variability in the TOD effect for the brain anatomy in a voxel– or vertex-wise manner.

## 2. Materials and Methods

### 2.1. Participants

77 volunteers were recruited for the study. All participants were right-handed, had normal or corrected to normal vision, no history of neurological and psychiatric disorders, and were drug-free. The following additional inclusion criteria were applied: age between 19 and 35 years, no shift work, not having been on a flight crossing more than two time zones within the past two months, declared regular daily schedule with no sleep debt (from 6.5 to 9 h of sleep each night). The level of daytime sleepiness was controlled with Epworth Sleepiness Scale (ESS; Johns 1991) and sleep quality with Pittsburgh Sleep Quality Index (PSQI; Buysse et al. 1989). The regularity of participants’ sleep-wake schedule was checked for one week prior to MRI scanning using MotionWatch8 actigraphs (CamNtech Ltd.). After inspection of the MRI data, five participants were excluded from the study due to unsatisfactory removal of meninges during the brain segmentation process. The final sample consisted of 72 participants (45 females). To ensure the validity of our results and estimate unbiased effect sizes, the preprocessed dataset was pseudorandomly split into two halves counterbalanced in regards to the used investigation protocols (see Figure 1), named primary and secondary cohorts. We attempted to replicate the findings observed in each half of the dataset in the remaining participants of the study. A summary of the demographics for the two datasets and the full sample is provided in Table 1.

**Figure 1.**
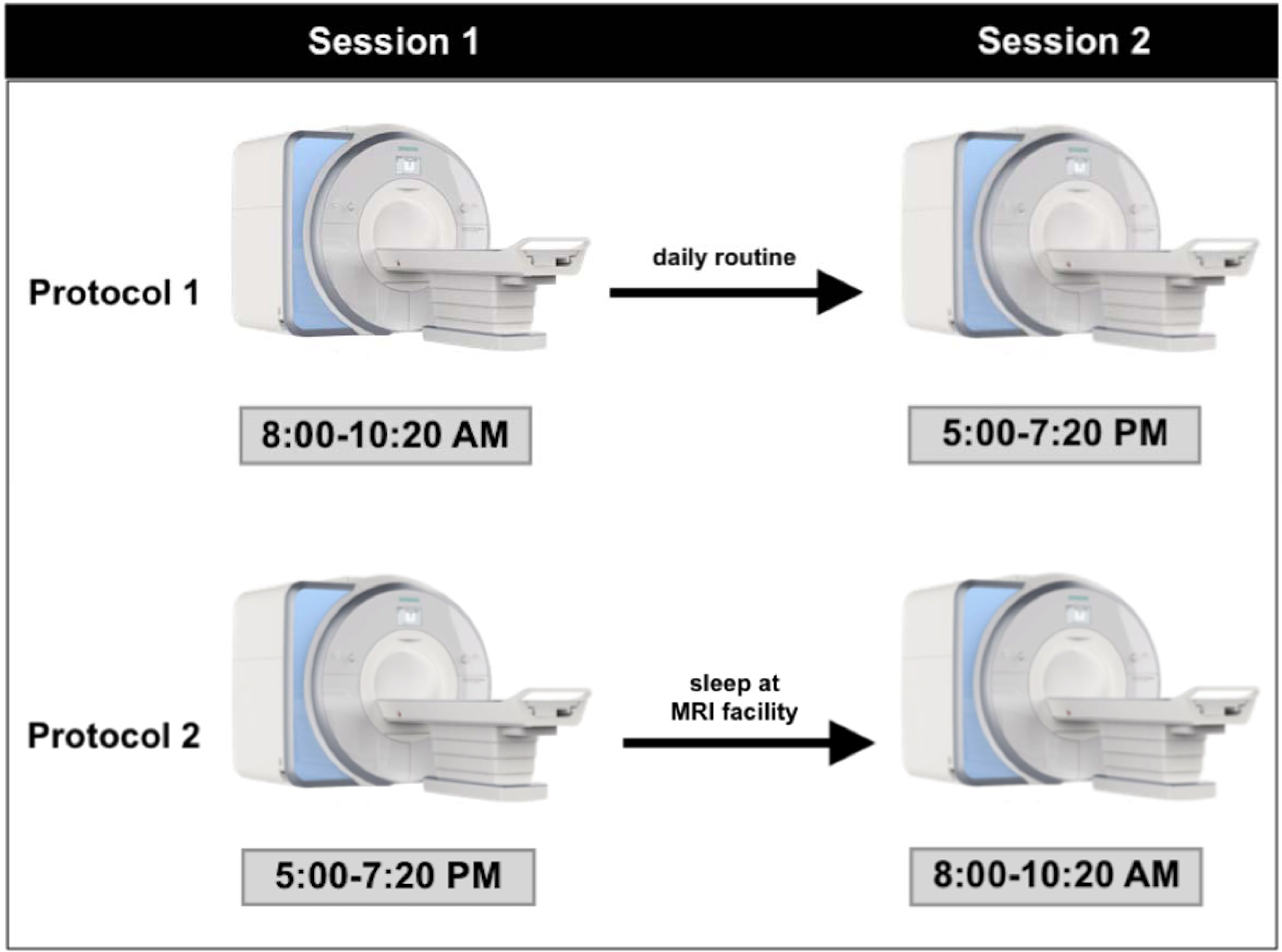
Visual demonstration of the study design. The number of participants undergoing protocol 1 and protocol 2 was counterbalanced across the primary and secondary datasets. In both protocols during the experimental days the participants were only allowed to engage in non-strenuous activities.

**Table 1.** Demographic and sleep characteristics of the sample.

| **Variable** | **Primary cohort, n = 36**<br>**(mean ± SD)** | **Secondary cohort, n = 36**<br>**(mean ± SD)** | **Group difference**<br>**(p value)** | **Full cohort, n = 72**<br>**(mean ± SD)** |
| --- | --- | --- | --- | --- |
| Sex (male/female) | 17/19 | 10/26 | NA | 27/45 |
| Age (years) | 23.78 ± 2.77 | 24.08 ± 3.77 | 0.841 <sup>NP</sup> | 23.93 ± 3.28 |
| Pittsburgh Sleep Quality Index | 2.86 ± 1.10 | 3.25 ± 1.05 | 0.177 <sup>NP</sup> | 3.06 ± 1.08 |
| Epworth Sleepiness Scale | 6.14 ± 2.68 | 6.47 ± 2.69 | 0.604 <sup>P</sup> | 6.30 ± 2.67 |
| Fall asleep time (hr) | 0:23 ± 1:10 <sup>M</sup> | 0:14 ± 1:00 | 0.560 <sup>P</sup> | 0:19 ± 1:05 <sup>M</sup> |
| Wake up time (hr) | 8:04 ± 1:15 <sup>M</sup> | 7:50 ± 1:18 | 0.454 <sup>P</sup> | 7:57 ± 1:17 <sup>M</sup> |
| Time in bed (hr) | 7:40 ± 0:51 <sup>M</sup> | 7:32 ± 0:43 | 0.449 <sup>P</sup> | 7:50 ± 0:49 <sup>M</sup> |
| Actual sleep time (hr) | 6:37 ± 0:44 <sup>M</sup> | 6:28 ± 0:36 | 0.329 <sup>P</sup> | 6:33 ± 0:41 <sup>M</sup> |
| Sleep efficiency<sup>1</sup> (%) | 84.22 ± 4.60 <sup>M</sup> | 83.39 ± 5.02 | 0.477 <sup>P</sup> | 83.78 ± 4.78 <sup>M</sup> |
| Sleep latency<sup>2</sup> (min) | 8.31 ± 7.42 <sup>M</sup> | 9.84 ± 11.76 | 0.968 <sup>NP</sup> | 9.13 ± 9.95 <sup>M</sup> |
| Sleep fragmentation index | 23.60 ± 6.08 <sup>M</sup> | 24.48 ± 6.56 | 0.566 <sup>P</sup> | 24.06 ± 6.31 <sup>M</sup> |

Abbreviations: SD = standard deviation; NA = non-applicable.
<sup>NP</sup> Assessed with a non-parametric Mann-Whitney U test.
<sup>P</sup> Assessed with a parametric two sample t-test.
<sup>M</sup> Actigraphic data missing for 3 participants.
<sup>1</sup> Sleep efficiency was calculated as the percentage of the actual sleep time divided by the time in bed.
<sup>2</sup> Time between switching off the lights and falling asleep.

The study protocol is presented in Figure 1. Each participant underwent two scanning sessions: one in the morning, and the other one in the evening, about 1 and 10 h after awakening, respectively. The beginning of experimental sessions (morning and evening) was adjusted to the declared and objective (derived from actigraphic data) sleep-wake patterns of participants on a regular working day. The morning scans were acquired between 08:00 and 10:20, whereas the evening scans were collected between 17:00 and 19:20. The session order was counterbalanced across participants in the full sample, as well as across the primary and secondary cohorts. Participants completed the study on one day (when the morning session was first) or on two consecutive days (when the evening session was first). On the night before the morning session participants slept in the rooms located in the same building as the MRI laboratory. They were only allowed to engage in non-strenuous activities during the experimental days. Participants abstained from alcohol (48 h) and caffeine (24 h) before the first session. Caffeine and alcohol were also banned during the study days. The study was conducted under the Declaration of Helsinki and approved by the Ethics Committee for Biomedical Research at the Military Institute of Aviation Medicine (Warsaw, Poland; 26 February 2013) and the Research Ethics Committee of the Institute of Applied Psychology at Jagiellonian University (Kraków, Poland; 21 February 2017).

As the current dataset was a part of a larger investigation into the effects of chronotype and subjective amplitude of diurnal variations in mood and cognition on brain morphometric measures (Zareba et al. 2021; Zareba et al., 2023), we additionally tested in the full sample whether there was any interaction between these circadian characteristics (Oginska 2011; Oginska et al. 2017) and TOD volumetric brain differences. No such effects were found (p > 0.05) and thus these measures were not included in the models.

### 2.2. Actigraphy-Derived Sleep Measurements

The sleep-wake patterns of the participants were monitored with MotionWatch8 actigraphs (CamNtech Ltd.). The devices were given to the subjects one week prior to MRI scanning; participants wore them non-stop on the wrist of the non-dominant hand. By pressing the ‘event’ button’, they marked the moments when the lights went out and when they woke up. The following parameters were extracted from full week registrations with use of the MotionWare Software: (1) average fall asleep time; (2) average wake-up time; (3) time in bed; (4) ‘actual’ or ‘total’ sleep time; (5) sleep efficiency (actual sleep time expressed as a percentage of time in bed); (6) sleep latency (time between switching off the lights and falling asleep); (7) sleep fragmentation index based on night-time movement data (higher values indicate greater mobility during the night – it can be used as an indication of sleep quality). Sleep length (actual sleep time) and sleep fragmentation index were chosen as the variables of interest for correlation with the replicated TOD findings. The complete actigraphic data from 3 participants was not available due to battery or connection failure, meaning the sleep parameters were available for 69 subjects. None of the sleep measures differed between the cohorts (p > 0.05). Their summary for both groups and the full sample is provided in Table 1.

### 2.3. MRI Data Acquisition

The high resolution, structural images used in the described investigation were acquired using a 3T scanner (Magnetom Skyra, Siemens) equipped with a 64-channel head/neck coil with a T1 MPRAGE sequence (field of view = 256 x 248 mm; 176 sagittal slices; 1×1×1.1 mm3 voxel size; repetition time = 2300 ms, echo time = 2.98 ms, inversion time = 900 ms, flip angle = 9°, GRAPPA acceleration factor 2). The anatomical MRI data was collected as a part of a longer protocol that additionally included other MRI modalities.

### 2.4. Voxel-Based Morphometry

The brain images underwent processing with the standard VBM pipeline (Ashburner and Friston 2000) in Computational Anatomy Toolbox (CAT12 version 12.8; run under MATLAB 2019b; Gaser et al., 2022). Firstly, each volume was segmented into GM, WM and CSF, which was followed by normalisation into the MNI152 space using diffeomorphic anatomical registration through exponentiated lie algebra (DARTEL). Such normalised GM and WM segments were subsequently modulated with the Jacobian determinant to account for the resulting signal intensity changes. The preprocessed GM and WM volumes were inspected visually to ensure satisfactory quality of the segmentation. Lastly, each of the segmented, normalised, and modulated images underwent smoothing with a 4-mm Gaussian filter.

The statistical analysis was performed separately for each tissue class using the 3dLME program (Chen et al. 2013) available in AFNI (Cox 1996). Voxels with estimated mean study-wise volumes lower than 0.1 were excluded from the analyses to prevent the inclusion of non-brain data. As mentioned before, the whole dataset of 72 participants was divided into two equal cohorts to attempt to replicate the findings observed in each half of the dataset and estimate their effect sizes in an unbiased manner using Cohen’s d. The models investigated the TOD effect using a paired t-test, with sex, age and total intracranial volume (ICV) controlled as covariates. The multiple comparison correction was achieved through cluster-level family-wise error (FWE) < 0.05 following the initial voxel-level thresholding (p < 0.001). The significant clusters from these GM and WM TOD models were saved separately as binary masks and were subsequently used in the replication analysis to extract the cluster-averaged brain tissue volumes in the other dataset. The statistical significance was assessed with paired t-tests, and the resulting p values were corrected for multiple comparisons using false discovery rate (FDR) to increase the replication power. As the VBM data can be affected by technical factors such as signal-to-noise ratio (SNR) or head motion (Goto et al. 2013; Pardoe et al. 2016; Pardoe and Martin 2022; Reuter et al. 2015), we furthermore calculated indices of data quality (see Supplementary Material) and ran in each cohort additional whole-brain models with data quality metrics serving as the variables of interest to ensure that the TOD effects did not overlap spatially with areas affected by these factors. The same covariates (sex, age and ICV) and statistical thresholds (cluster level FWE < 0.05 following voxel-level threshold of p < 0.001) as in the TOD analyses were used. The results are presented in Supplementary Material Tables 5-6 and Supplementary Material Figures 7-8.

## 3. Results

### 3.1. Time-of-Day Differences in Voxel-Based Morphometry in Primary Dataset

The TOD differences from the whole-brain analysis in the primary dataset are depicted in Table 2 and Figure 2. The evening scans were associated with decreased WM volume in the bilateral parietal lobes and increased WM volume in the right internal capsule. Furthermore, the evening session was characterised by lower GM volume in the bilateral retrosplenial cortex, together with the right occipital pole, inferior temporal gyrus, thalamus and cerebellar lobule VIIa. The clusters of increased WM volume in the right internal capsule and decreased GM volume in the right thalamus largely overlapped with each other (see Figure 2). However, these two areas were also affected by data quality in the primary cohort (see Supplementary Material Figure 7 and Supplementary Material Table 5). No other regions indicated in the primary TOD analysis were found to overlap with the data quality results in any of the two datasets.

**Figure 2.**
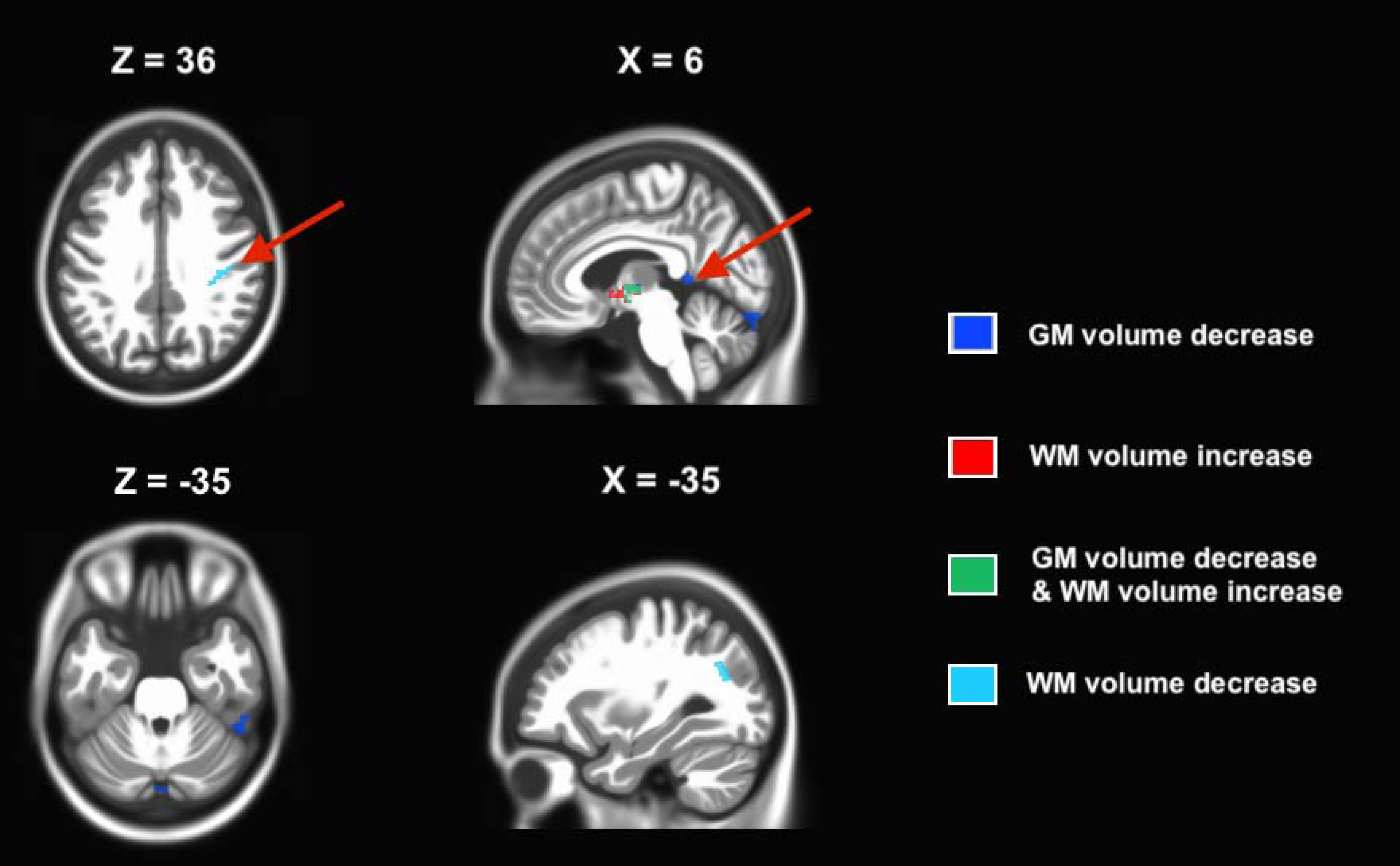
The time-of-day (TOD) effect on the voxel-based morphometry (VBM) in the primary dataset (cluster-level FWE < 0.05). Red arrows indicate clusters that were also significant in the secondary cohort. Abbreviations: GM, grey matter; WM, white matter.

**Table 2.**
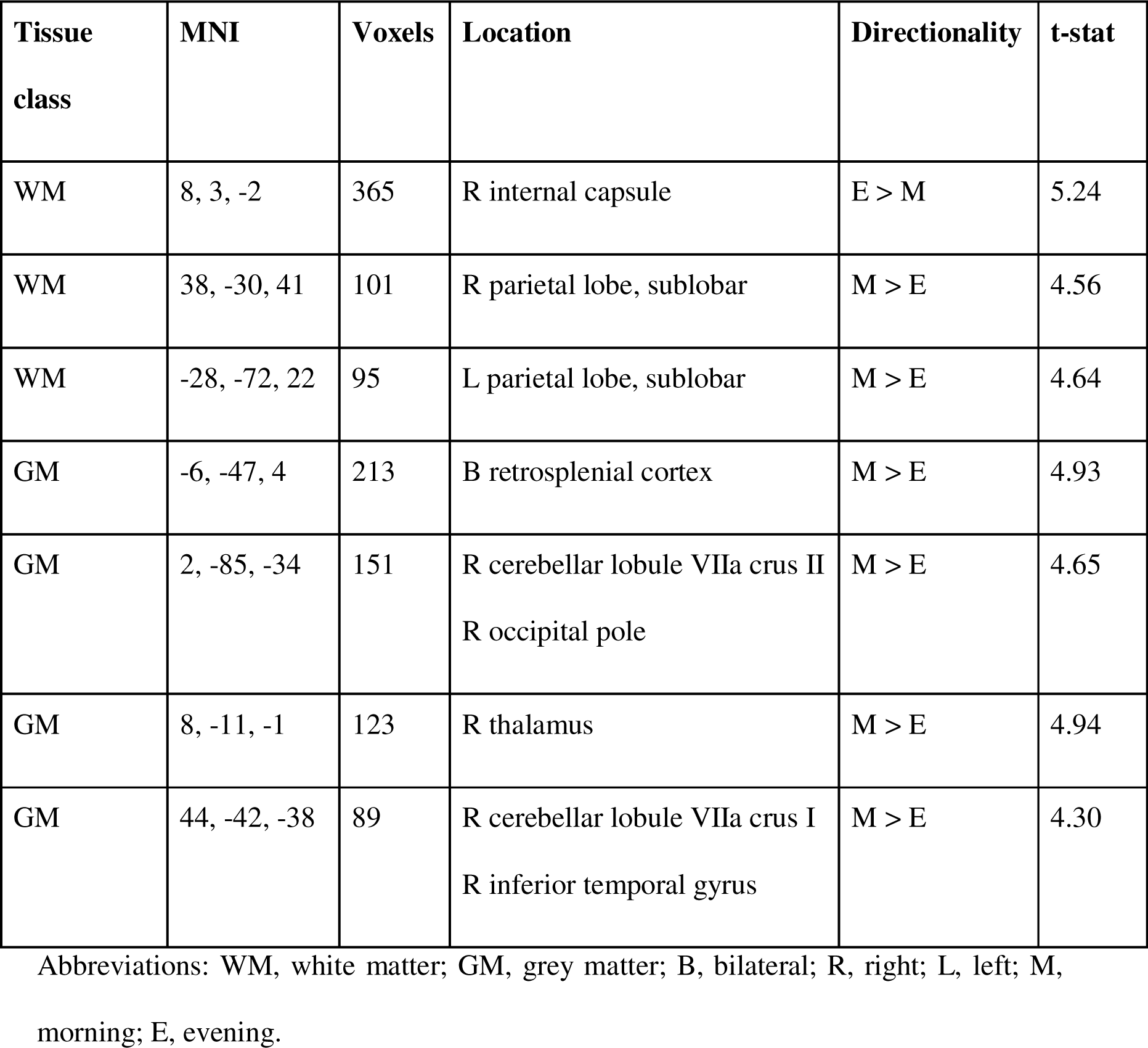
Time-of-day (TOD) differences in the voxel-based morphometry (VBM) in the primary dataset (cluster-level FWE < 0.05).

The comparisons in the second dataset replicated with medium effect sizes 2 clusters from the primary investigation, i.e. morning-to-evening decreases in GM volume in the bilateral retrosplenial cortex (Cohen’s d = –0.633; FDR = 0.004) and WM volume in the right parietal lobe (Cohen’s d = –0.568; FDR = 0.006). The full results of the replication analysis of the findings from the primary cohort are presented in Supplementary Material Table 7.

### 3.2. Time-of-Day Differences in Voxel-Based Morphometry in Secondary Dataset

The TOD differences from the whole-brain analysis in the secondary dataset are shown in Table 3 and Figure 3. The evening scans were characterised by decreased WM volume in the bilateral frontal, temporal and parietal lobes, as well as the left occipital regions. The only area of increased WM volume was located in the brainstem. The evening session was further associated with increased GM volume in the bilateral frontal, temporal, occipital and cerebellar cortex. Additionally, larger GM volumes were found in the left amygdala and hippocampus, as well as the right angular gyrus. The only TOD decrease in GM volume was located in the bilateral anterior cingulate cortex. The clusters of increased GM volume and decreased WM volume were found to partially overlap in the left orbitofrontal cortex, amygdala and hippocampus, fusiform and inferior occipital gyri, as well as the right angular gyrus and temporoparietal junction. None of the results found in the secondary cohort mapped onto regions affected by data quality in both datasets, with the exception of the left amygdala and hippocampus together with the adjacent WM, which was affected by this factor solely in the secondary dataset (see Supplementary Material Figure 8 and Table 6).

**Figure 3.**
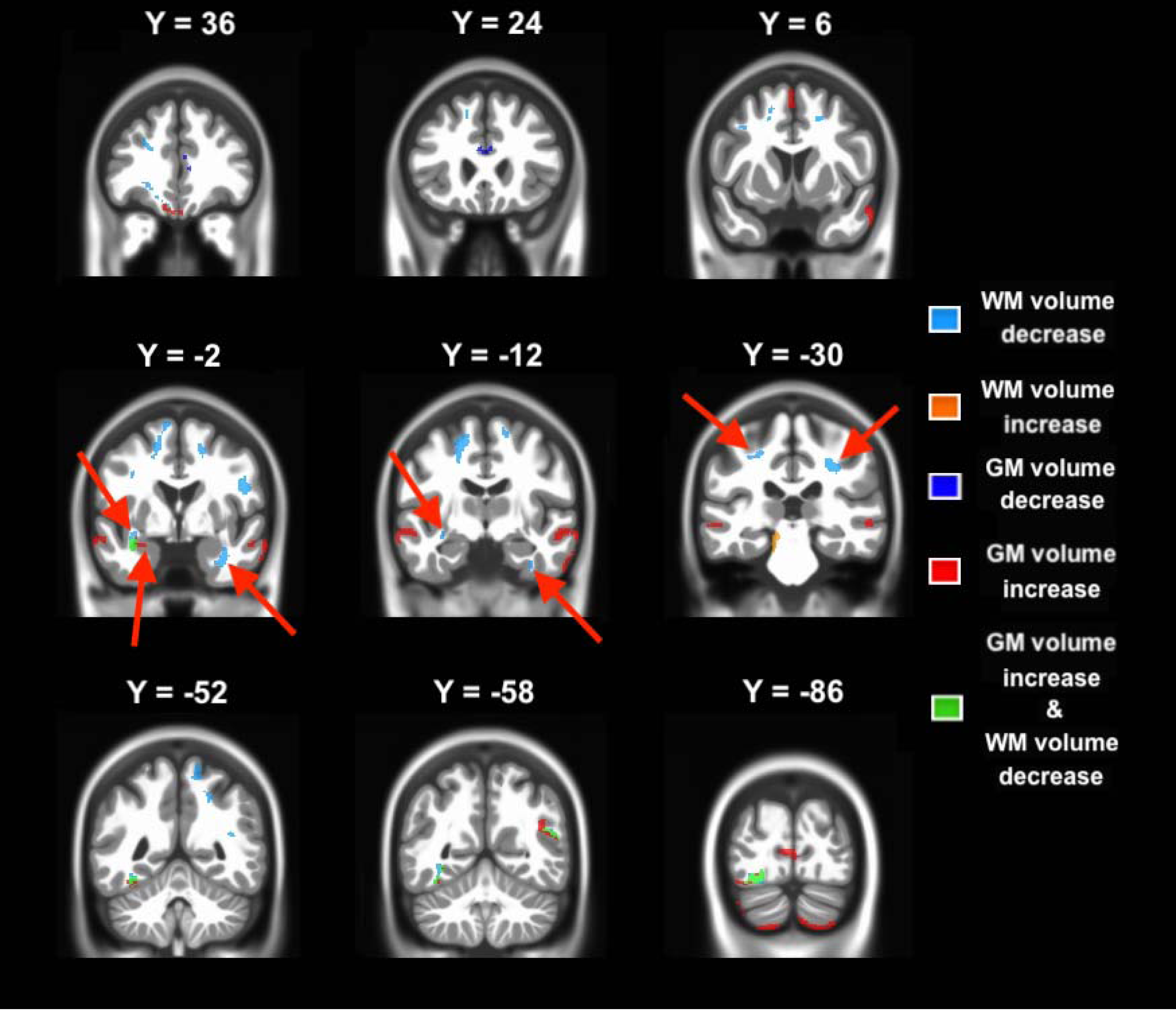
The time-of-day (TOD) effect on the voxel-based morphometry (VBM) in the secondary dataset (cluster-level FWE < 0.05). Red arrows indicate clusters that were also significant in the primary cohort. Abbreviations: GM, grey matter; WM, white matter.

**Figure 4.**
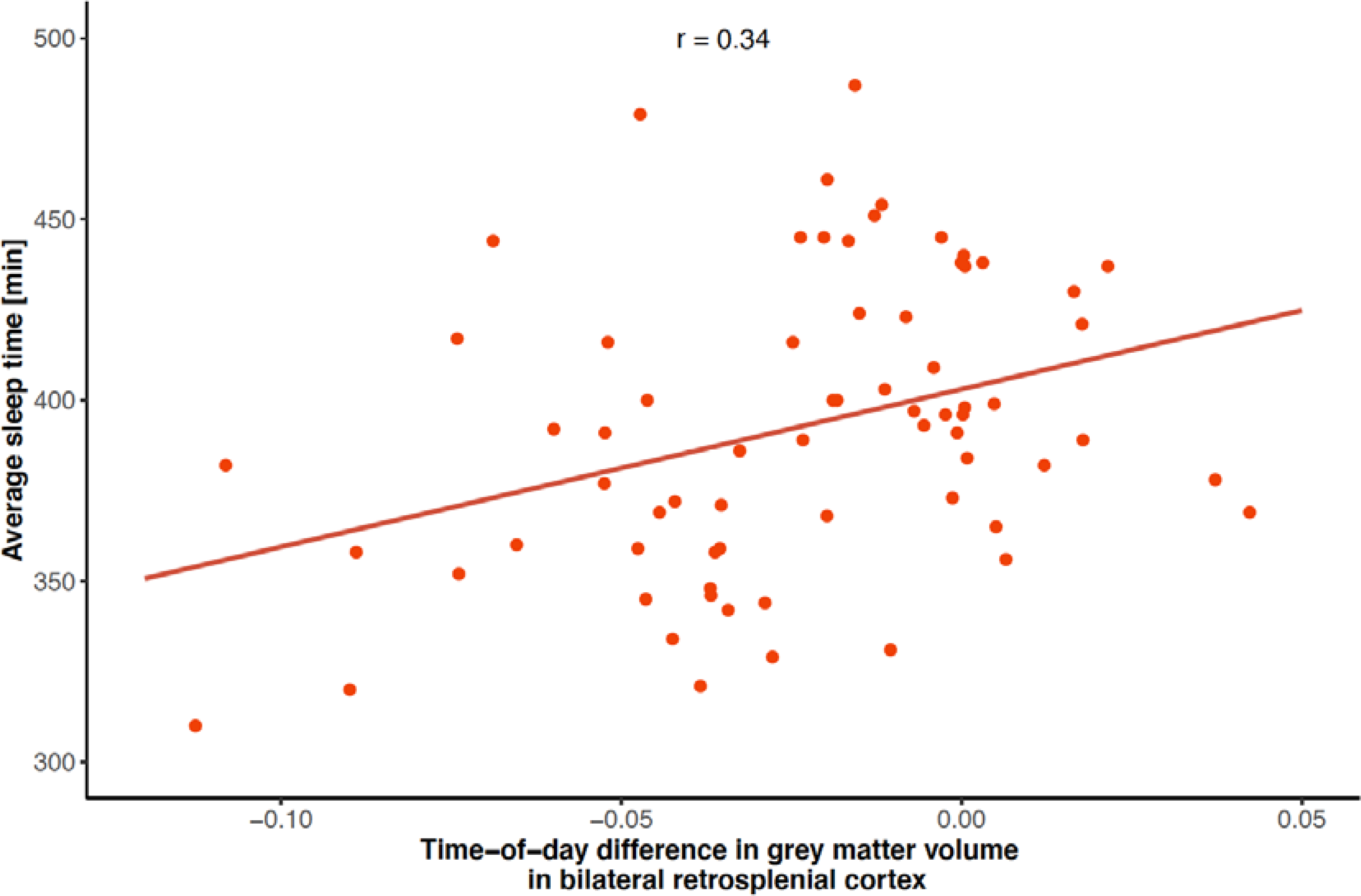
The TOD change in the GM volume in the bilateral retrosplenial cortex was positively correlated with the average sleep time over the course of the previous week (r = 0.34).

**Table 3.** Time-of-day (TOD) differences in the voxel-based morphometry (VBM) in the secondary dataset (cluster-level FWE < 0.05).

| **Tissue class** | **MNI** | **Voxels** | **Location** | **Directionality** | **t-stat** |
| --- | --- | --- | --- | --- | --- |
| WM | -13, -5, 60 | 697 | L parietal, subgyral | M > E | 5.63 |
| WM | 26, -35, 45 | 630 | R parietal, sublobar | M > E | 6.27 |
| WM | -34, -61, -11 | 375 | L occipital, subgyral | M > E | 5.66 |
| WM | -31, -8, -15 | 226 | L temporal, sublobar | M > E | 6.29 |
| WM | 18, 7, 49 | 187 | R parietal, subgyral | M > E | 5.61 |
| WM | 33, 0, -38 | 174 | R temporal, sublobar | M > E | 5.50 |
| WM | 47, -4, 29 | 172 | R parietal, subgyral | M > E | 5.79 |
| WM | -37, -80, -22 | 152 | L occipital, subgyral | M > E | 4.88 |
| WM | -10, -32, -19 | 143 | Brainstem | E > M | 5.18 |
| WM | -30, -29, 49 | 136 | L parietal, sublobar | M > E | 5.27 |
| WM | -16, 46, 26 | 134 | L frontal, sublobar | M > E | 5.02 |
| WM | -22, -63, 43 | 134 | L parietal, subgyral | M > E | 4.28 |
| WM | 15, -11, 68 | 134 | R frontal, subgyral | M > E | 5.16 |
| WM | 11, -56, 66 | 127 | R parietal, subgyral | M > E | 4.78 |
| WM | -21, -39, 55 | 124 | L parietal, sublobar | M > E | 4.71 |
| WM | -18, 32, -16 | 123 | L orbitofrontal | M > E | 4.34 |
| WM | -28, -44, 37 | 112 | L parietal, sublobar | M > E | 4.53 |
| WM | 39, -50, 17 | 110 | R temporoparietal junction | M > E | 4.51 |
| WM | 17, -5, 53 | 101 | R frontal, subgyral | M > E | 5.53 |
| WM | -19, 41, -7 | 97 | L orbitofrontal | M > E | 4.41 |
| WM | -30, -1, 36 | 96 | L frontal, subgyral | M > E | 4.21 |
| GM | 61, -6, -15 | 1003 | R middle temporal gyrus | E > M | 5.40 |
| GM | 0, 63, 0 | 720 | B frontal pole<br>L orbitofrontal cortex | E > M | 6.32 |
| GM | -62, -24, -9 | 572 | L middle temporal gyrus | E > M | 8.02 |
| GM | 15, -76, -71 | 310 | R cerebellar cortex | E > M | 4.96 |
| GM | -37, -80, -22 | 215 | L inferior occipital gyrus | E > M | 5.35 |
| GM | -33, -6, -15 | 143 | L amygdala<br>L hippocampus | E > M | 5.01 |
| GM | -15, -83, -64 | 128 | L cerebellar cortex | E > M | 4.67 |
| GM | 8, 30, 19 | 119 | B anterior cingulate cortex | M > E | 4.96 |
| GM | -30, -64, -11 | 118 | L fusiform gyrus | E > M | 4.67 |
| GM | -34, -54, -21 | 116 | L fusiform gyrus | E > M | 4.61 |
| GM | 58, -31, -5 | 105 | R middle temporal gyrus | E > M | 5.46 |
| GM | 0, 6, 70 | 105 | B superior frontal gyrus | E > M | 5.51 |
| GM | 42, -59, 27 | 103 | R angular gyrus | E > M | 4.73 |
| GM | -39, -80, -57 | 98 | L cerebellar cortex | E > M | 4.06 |
| GM | -10, -89, 2 | 98 | B cuneus | E > M | 4.99 |

Abbreviations: WM, white matter; GM, grey matter; B, bilateral; R, right; L, left; M, morning; E, evening.

The analysis in the primary dataset successfully replicated with medium effect sizes 5 clusters, i.e. morning-to-evening increases in GM volume in the left amygdala and hippocampus (Cohen’s d = 0.531; FDR = 0.035), as well as bilaterally symmetrical TOD decreases in WM volume in adjacent temporal lobes (Coheńs d = –0.530 and – 0.641, FDR = 0.035 and 0.018, for the left– and right-sided region respectively) and sublobar parietal WM (Coheńs d = –0.515 and –0.495, FDR = 0.035 and 0.038, for the left– and right-sided cluster respectively). The full description of the replication analysis in the primary cohort is presented in the Supplementary Material Table 8.

### 3.3. Association between Time-of-Day Results and Sleep Indices

Sleep time, sleep fragmentation and the replicated TOD volumetric differences were all normally distributed according to the Kolmogorov–Smirnov test (p > 0.05), and thus Pearson’s correlations were calculated to evaluate the associations between sleep indices and structural brain differences. As there was a substantial overlap between the decreases in the WM volume in the right parietal lobe (see Supplementary Material Figure 9), the clusters from the two analyses were merged. The results are presented in Table 4. The only significant correlation was found between the TOD GM effect in the bilateral retrosplenial cortex and sleep time (r = 0.34, FDR = 0.029; see Figure 2). At the nominal level, there was also a negative relation between the sleep fragmentation index and TOD volumetric difference in the right temporal WM adjacent to amygdala and hippocampus (r = –0.28, p = 0.02). However, as stated before, this finding did not survive the correction for multiple comparisons and is outlined here in case future studies with higher statistical power may want to investigate this matter more closely.

**Table 4.** Correlations between the actigraphy-derived indices of sleep and regional time-of-day (TOD) differences in grey matter (GM) and white matter (WM) volume. Bolded values signify successful replication, while italics indicate uncorrected p values < 0.05.

| **Tissue class** | **MNI** | **Voxels** | **Location** | **Sleep time correlation (FDR)** | **Sleep fragmentation correlation (FDR)** |
| --- | --- | --- | --- | --- | --- |
| GM | -6, -47, 4 | 213 | B retrosplenial cortex | **0.34 (0.029)** | -0.18 (0.456) |
| GM | -33, -6, -15 | 143 | L amygdala<br>L hippocampus | 0.12 (0.626) | 0.01 (0.993) |
| WM | 26, -35, 45 | 691 | R parietal, sublobar | 0.09 (0.626) | -0.05 (0.939) |
| WM | -31, -8, -15 | 226 | L temporal, sublobar | -0.17 (0.56) | -0.01 (0.993) |
| WM | 33, 0, -38 | 174 | R temporal, sublobar | 0.10 (0.626) | -0.28 (*0.138*) |
| WM | -30, -29, 49 | 136 | L parietal, sublobar | 0.04 (0.861) | -0.13 (0.644) |

Abbreviations: FDR, false discovery rate.

## 4. Discussion

To the best of our knowledge, our work is the first one to simultaneously consider the TOD effect on WM and GM VBM. The study advances the field of circadian neuroscience by showing that VBM-measured WM volume is significantly affected by TOD, with a vast majority of regions undergoing volumetric decreases. Additionally, we broaden the state of knowledge by providing evidence for TOD-related increases in GM volume, extending the previous findings of solely negative associations (Trefler et al., 2016). Last but not least, our analysis also reveals for the first time that the magnitude of structural TOD differences is related to individual sleep health.

### 4.1. Influence of Time-of-Day on Voxel-Based Morphometry

After excluding the results overlapping with the data quality-related findings, predominantly morning-to-evening reductions in WM volume were observed across two datasets. Thus, we add onto the literature which shows that TOD affects the diffusion of water molecules within WM tracts (Jiang et al. 2014; Thomas et al. 2018; Voldsbekk et al. 2020). Distinguishing between VBM and diffusion tensor imaging-based metrics is important especially that the results obtained with these two methods do not always converge, suggesting that the two methodologies differ in their sensitivity to certain aspects of WM microstructure (Padovani et al. 2005; Yoon et al. 2011). The uniform character of WM volumetric decreases observed across the two datasets, as well as their higher replicability than in the case of GM results (6 significant WM clusters at the corrected level, 7 more showing trend-level replicability) suggests that in the TOD context typical for MRI studies the effects associated with increased CSF-like water diffusivity might have greater impact on the WM signal characteristics than the experience-dependent plasticity. This is in line with previous reports stating that the amount of fluid detectable in perivascular spaces of WM increases with TOD (Thomas et al., 2018; Barisano et al. 2021), while oligodendrocyte proliferation, a key process for myelination and thus WM development, is more pronounced during sleep (Li et al., 2017).

In the case of GM, after accounting for the data quality, we observed opposite effects in the primary and secondary datasets, i.e. volumetric decreases and increases, respectively. The only study probing the TOD effect for GM reported that 4 hours of controlled rest were associated with lower GM volumes (Trefler et al., 2016), which was linked to the increased amount of CSF-like water in the superficial cortical layers by a follow-up work (Thomas et al., 2018; Xie et al., 2013). Nevertheless, as stated in the introduction, we believed that the experience-dependent plasticity could also take place if participants were allowed to engage in their usual, non-strenuous activities during a typical day (Hübener and Bonhoeffer 2010). Indeed, this alteration in the experimental design, together with increasing the differences in circadian times of data acquisition from 4 to 9 hours, has led to widespread increases in GM volume across multiple brain areas in the secondary dataset, resembling the effects of early sleep deprivation on brain anatomy (Dai et al., 2018).

However, as mentioned before, these effects were present only in one of the two cohorts. Indeed, we observed a high variability of GM TOD effects, with only two successfully replicated clusters. The first of them was located in the bilateral retrosplenial cortex, bordering the CSF compartment, and exhibited volumetric decrease in both datasets, which aligns well with the previously observed results (Thomas et al., 2018: Trefler et al., 2016). Interestingly, the second successfully replicated effect was situated in the left amygdala and hippocampus, two areas playing a fundamental role in emotional and episodic memory (Packard et al., 1989; Squire and Zola-Morgan, 1991). One fMRI study deploying a comparable experimental design to that of our own found a pronounced morning to evening increase in resting-state functional connectivity between the medial temporal lobes and other brain areas involved in memory functions (Shannon et al., 2013). Within that context, our results may be, at least partially, related to memory formation. Furthermore, two of the replicated WM clusters were found bilaterally in the parts of temporal lobes adjacent to the amygdalo-hippocampal area, with the left hemispheric WM volume decrease partially overlapping with areas of increased GM volume. Even though these areas were affected by head motion in the secondary dataset, their successful replication in the primary cohort indicates that it is indeed a physiological effect. This suggests that in the voxels located at the border of these two tissues, the synaptic plasticity may be one of the driving forces for the observed volumetric GM increase, a metabolically demanding process that may increase locally the homeostatic sleep pressure, which can manifest itself as elevated permeability of CSF-like water into the brain parenchyma (Thomas et al., 2018) and thus WM volume decrease. Similar processes may also be contributing to the volumetric WM effects across the other brain areas, however, we cannot exclude the presence of other region-specific mechanisms, especially that the spatial overlap between WM and GM findings was observed only in a handful of anatomical locations.

Regardless of the variability in the results, our study makes a major step forward by estimating unbiased effect sizes of the TOD effects. The replicated clusters from both analyses indicated medium effect strength, which underlines the importance of accounting for the TOD in neuroimaging studies. However, tightly controlled conditions are often difficult to arrange in the research settings. Thus, we suggest that limiting the impact of TOD on VBM should be achieved through at least scheduling participants’ scans to a narrow window of time since their (habitual) wake up times.

### 4.2. Association between Sleep and Daily Differences in Voxel-Based Morphometry

The daily change in the GM volume in the bilateral retrosplenial cortex had a moderate correlation with the average sleep time of the participants during the previous week (r = 0.34). This finding serves as a further premise for the biological basis of the observed TOD results, and suggests that shorter sleep may contribute to higher cortical permeability for the CSF-like water pool, a process that could be related to insufficient sleep pressure dissipation. Furthermore, it expands the existing literature which shows that the sleep-related factors modulate the relationship between circadian rhythms and brain activity (Song et al. 2019; Vandewalle et al. 2011).

Contrary to our hypothesis, the relationship between GM volume differences in the bilateral retrosplenial cortex and sleep length was the only significant association between the daily fluctuations in tissue volume and measures of sleep health. It cannot be excluded that variation in sleep length and sleep fragmentation exhibits a stronger effect on the diffusivity of CSF-like water pool in general rather than the TOD fluctuation in this metric. Poor sleep quality has been associated with enlarged perivascular spaces and increased CSF tracer enrichment (Berezuk et al. 2015; Eide et al. 2022). While the amount of fluid detectable within perivascular spaces increases with TOD (Barisano et al. 2021) and has been shown to be under circadian control (Hablitz et al. 2020), no studies have examined the impact of sleep health on the TOD differences in the discussed measures. Future studies are thus warranted to fill this crucial gap in our understanding of the described phenomena.

### 4.3. Limitations

As discussed before, we observed a high variability of the GM results between the two datasets. Although the participants abstained from alcohol and caffeine consumption prior to the experiment, an additional, uncontrolled source of the between-subject variability in the TOD effects could come from factors that are believed to impact the glymphathic functions, such as hypertension and hydration levels (Mortensen et al., 2019; Yi et al., 2022), with the latter additionally directly linked to alterations in GM and WM volumes (Streitbürger et al., 2012; Zhang et al., 2022). Another potential explanation for the low replicability of the GM TOD differences could be the lack of strict control of between-session activity of the participants, aimed at mimicking the designs of a vast majority of MRI studies. As a result, our findings represent a combination of circadian– and experience-dependent effects, and it cannot be excluded that the exact nature of the cognitive processes the participants engaged in throughout the day could have altered the regional balance between the plasticity and CSF-permeability related processes, and as such impact the measurements. Thus, it would be interesting to investigate how altered the results would be if the subjects instead engaged in a cognitive training that affects only specific neural networks, for example by building on some earlier diffusion studies in humans and animals (Hofstetter et al. 2013). Such an approach could also enable to study the associations of the structural brain differences with homeostatic sleep pressure, a process that has been linked to the brain activity during wakefulness (Hung et al., 2013) and has been suggested to modulate the permeability of CSF-like water into the brain parenchyma (Thomas et al., 2018). Nevertheless, the present experimental design can also be considered one of the major strengths of our work as neuroimaging studies often do not account for the time of data acquisition and participants’ activities performed on the day of the scan prior to the visit in the laboratory, thus making our findings a representative of the effects that could be otherwise misattributed to experimental manipulation.

On a further note, as our datasets were collected only in young individuals and ageing has been shown to impact the glymphatic functions and synaptic plasticity (Bloss et al. 2011; Salman et al. 2022), we believe that the results presented in this manuscript may not be fully replicable in the older groups. Last but not least, despite our interest in investigating the relationship between the sleep data and TOD variability in WM and GM volume, we restricted the currently described analyses only to the individuals without any sleep-related disorders. Future studies are therefore warranted to enrich our knowledge by responding to these limitations.

## 5. Conclusions

Our work extends the field of circadian neuroscience by reporting that TOD is predominantly associated with medium effect size decreases in regional WM volume, and provides the first evidence for volumetric GM increases of comparable strength across the same time frame, stressing out the need to control for TOD in structural neuroimaging research. The WM effects, likely related to the permeability of CSF-like water pool and homeostatic sleep pressure, were characterised by higher replicability than structural GM differences. Across the two datasets, we observed GM volumetric increases in amygdala and hippocampus, and decreases in retrosplenial cortex, with the latter being more pronounced in individuals with shorter sleep times. These findings implicate the potential existence of region-specific mechanisms driving the GM effects, which might be modulated by cognitive processes taking place during wakefulness and homeostatic sleep pressure.

## Data availability

The dataset used in this study will be made available upon manuscript acceptance.

## CRediT author statement

**MRZ:** Conceptualization, Methodology, Investigation, Formal Analysis, Validation, Data Curation, Writing – Original Draft, Visualization. **MF:** Conceptualisation, Methodology, Investigation, Data Curation, Writing – Review & Editing, Funding Acquisition. **TM:** Conceptualisation, Methodology, Writing – Review & Editing, Funding Acquisition. **HO:** Investigation, Formal Analysis, Writing – Review & Editing. **EB:** Investigation, Formal Analysis, Validation, Writing – Review & Editing. **AD:** Conceptualisation, Methodology, Investigation, Formal Analysis, Validation, Data Curation, Writing – Review & Editing, Supervision. All authors contributed to the article and approved the submitted version.

## Acknowledgements

This research was supported in part by PL-Grid Infrastructure. We thank Prof. Patricia Reuter-Lorenz for her constructive suggestions during the planning and development of the Harmonia project and her valuable support. We also thank Anna Beres, Piotr Faba, Koryna Lewandowska, Monika Ostrogorska, Barbara Sikora-Wachowicz for their assistance with the fMRI data collection, Aleksandra Zyrkowska for help with the process of participant selection, and Magdalena Debowska for help in collecting actigraphy data.

## Ethical statement

The authors declare that all experiments on human subjects were conducted in accordance with the Declaration of Helsinki and that all procedures were carried out with the adequate understanding and written consent of the subjects. The study was approved by the Ethics Committee of the Institute of Applied Psychology at Jagiellonian University (approval code: 3/2017).

