## Supplementary Materials for "Tracing Diurnal Differences in Brain Anatomy with Voxel-Based Morphometry – Associations with Sleep Characteristics"

**MRI Data Quality Assessment**

All the images that entered the analyses were found to have satisfactory (n = 19) or good data quality (n = 125) according to the quality check reports generated by the Computational Anatomy Toolbox (CAT12 version 12.8; run under MATLAB 2019b). As the next step, we applied the Preprocessed Connectomes Project Quality Assessment Protocol (PCP QAP; Shehzad et al. 2015) to further investigate the matter. For each T1-weighted scan, six quality metrics were calculated: signal-to-noise ratio (SNR; mean GM intensity divided by the standard deviation (SD) of background voxels), contrast-to-noise ratio (CNR; the difference between the mean WM and GM intensities divided by the SD of background voxels), foreground-to-background energy ratio (FBER; variance of voxels inside of head divided by variance of voxels outside of it), voxel smoothness as full width at half maximum (FWHM), entropy focus criterion (EFC; Shannon entropy of voxel signal in proportion to the highest possible entropy for a similarly sized image; it indicates motion and ghosting artefacts) and fraction of artefact background voxels (QI1; the proportion of background voxels with higher signal intensities than the most frequently occuring value). Their distributions across the dataset are shown in the Supplementary Material Figures 1-6.

The within-subject comparisons revealed significant between-session differences in the image quality for all the metrics but FWHM (see Supplementary Material Table 1 for more details). As moderate and strong correlations were found between the quality indices (see Supplementary Material Table 2), principal component analysis (PCA) was performed in R (princomp function, version 4.0.3) to reduce the data dimensionality and identify the factors most contributing to scan quality variability (see Supplementary Material Tables 3-4 for results). In line with the Kaiser criterion (i.e. component eigenvalues higher than 1), the first three components, accounting together for 90.7% of the variance, were chosen for further analyses. The first component (PC1) was most strongly associated with positive loadings from SNR, CNR and FBER, the second component (PC2) was most strongly related to negative loadings from EFC and QI1, whereas the third component (PC3) was mostly associated with a positive loading from FWHM. PC1 was found to have lower values for the evening session scans (paired t-test, T-stat = -6.06, p < 0.001), the opposite was observed for PC2 (paired t-test, T-stat = 5.08, p < 0.001) and no between-session differences were found for PC3 (paired t-test, T-stat = 0.89, p = 0.376). Thus, the models testing the influence of data quality on voxel-based morphometry (VBM) metrics included PC1 and PC2 as the variables of interest instead of the time-of-day (TOD).

The two cohorts did not differ in terms of the PC1 values in both sessions, nor in the PC2 values for the morning session (p > 0.05). A significant difference was observed for the PC2 in the evening session (two sample t-test, T-stat = 2.82, family-wise error rate (FWE)-corrected p = 0.037) with the secondary cohort displaying higher values (lower data quality) compared to the primary cohort. Nevertheless, the PC2 in the secondary cohort spanned across a broader range of values (from -3.02 to 1.92) than in the primary cohort (from -1.80 to 2.42). Additionally, the two groups had similar between-session differences in both PC1 and PC2 values (p > 0.05). As such, we believe that this factor has a limited impact on the results of the subsequently described analyses.

**Supplementary Material Table 1.** Between-session differences in the image quality metrics.

| Image quality metric | Mean^a^/Median^b^ | SD^a^/MAD^b^ | T-stat | p _uncorrected_ | Evening session data quality |
| --- | --- | --- | --- | --- | --- |
| SNR | -3.56^a^ | 5.21^a^ | 5.78 | < 0.001 | lower |
| CNR | -1.57^a^ | 2.43^a^ | 5.46 | < 0.001 | lower |
| FBER | -67.72^a^ | 105.38^a^ | 5.45 | < 0.001 | lower |
| FWHM | 0.02^a^ | 0.35^a^ | 0.61 | 0.543 | no difference |
| EFC | -0.03^b^ | 0.04^b^ | n/a | < 0.001 | higher |
| QI1 | -0.02^a^ | 0.06^a^ | 3.45 | < 0.001 | higher |

^a^ Means and standard deviations (SDs) indicate the use of the paired t-test. ^b^ Medians and median absolute deviations (MADs) indicate the use of the Wilcoxon test. Abbreviations: SNR, signal-to-noise ratio; CNR, contrast-to-noise ratio; FBER, foreground-to-background energy ratio; FWHM, full width at half maximum; EFC, entropy focus criterion; QI1, fraction of artefact background voxels; n/a, non-applicable.

**Supplementary Material Table 2.** Pearson’s correlation between image quality metrics across both sessions.

| Quality metric | SNR | CNR | FBER | FWHM | EFC | QI1 |
| --- | --- | --- | --- | --- | --- | --- |
| SNR | 1.00 |  |  |  |  |  |
| CNR | 0.98 | 1.00 |  |  |  |  |
| FBER | 0.86 | 0.84 | 1.00 |  |  |  |
| FWHM | -0.02 | -0.08 | -0.08 | 1.00 |  |  |
| EFC | 0.25 | 0.26 | 0.10 | -0.08 | 1.00 |  |
| QI1 | -0.09 | -0.10 | -0.09 | 0.16 | 0.56 | 1.00 |

Abbreviations: SNR, signal-to-noise ratio; CNR, contrast-to-noise ratio; FBER, foreground-to-background energy ratio; FWHM, full width at half maximum; EFC, entropy focus criterion; QI1, fraction of artefact background voxels.

**Supplementary Material Table 3.** Components derived from the principal component analysis (PCA).

| Component no. | 1^*^ | 2^*^ | 3^*^ | 4 | 5 | 6 |
| --- | --- | --- | --- | --- | --- | --- |
| SD | 1.692 | 1.250 | 1.011 | 0.614 | 0.394 | 0.138 |
| Proportion of explained variance | 0.477 | 0.260 | 0.170 | 0.062 | 0.025 | 0.006 |
| Cumulative explained variance | 0.477 | 0.737 | 0.907 | 0.969 | 0.994 | 1 |

The asterisks indicate components chosen for further analyses. Abbreviations: SD, standard deviation.

**Supplementary Material Table 4.** Loadings of the original image quality metrics on the first three principal components.

| Quality metric | Component 1 | Component 2 | Component 3 |
| --- | --- | --- | --- |
| SNR | 0.579 | 0.030 | 0.090 |
| CNR | 0.577 | 0.030 | 0.039 |
| FBER | 0.540 | 0.098 | 0.070 |
| FWHM | -0.066 | -0.136 | 0.958 |
| EFC | 0.185 | -0.662 | -0.250 |
| QI1 | -0.035 | -0.729 | 0.064 |

Abbreviations: SNR, signal-to-noise ratio; CNR, contrast-to-noise ratio; FBER, foreground-to-background energy ratio; FWHM, full width at half maximum; EFC, entropy focus criterion; QI1, fraction of artefact background voxels.

**Supplementary Material Table 5.** Data quality-related differences in VBM in the primary dataset (cluster-level FWE < 0.05).

| **Tissue class** | **MNI** | **Voxels** | **Location** | **Component** | **F-stat** |
| --- | --- | --- | --- | --- | --- |
| WM | 12, -6, -3 | 291 | R internal capsule | PC1 | 37.00 |
| WM | 15, 51, -14 | 178 | R orbitofrontal cortex | PC1 | 32.06 |
| WM | -1, -2, -2 | 103 | L internal capsule  L fornix | PC1 | 36.25 |
| WM | -13, -82, -15 | 176 | L occipital lobe, subgyral | PC2 | 30.28 |
| GM | 42, -2, 7 | 274 | R insula | PC1 | 50.14 |
| GM | 2, 23, 31 | 224 | B anterior cingulate cortex | PC1 | 27.94 |
| GM | 6, -14, -1 | 157 | R thalamus | PC1 | 28.47 |
| GM | 64, 8, 28 | 146 | R precentral gyrus | PC1 | 21.86 |
| GM | 47, -5, -15 | 133 | R superior temporal gyrus | PC1 | 26.20 |
| GM | -43, -73, -28 | 123 | L cerebellar lobule VIIa crus I  L inferior occipital gyrus | PC1 | 26.99 |
| GM | 47, -73, 36 | 109 | R angular gyrus | PC1 | 20.02 |
| GM | -58, -43, 21 | 105 | L superior temporal gyrus | PC1 | 26.60 |
| GM | -4, -68, 68 | 105 | L precuneus | PC1 | 56.55 |
| GM | -19, -30, 77 | 105 | L postcentral gyrus | PC1 | 34.48 |
| GM | -18, -82, -21 | 114 | L lingual gyrus | PC2 | 36.26 |
| GM | 21, -100, 15 | 105 | R superior occipital gyrus | PC2 | 36.60 |

Abbreviations: WM, white matter; GM, grey matter; B, bilateral; R, right; L, left; PC1, principal component 1; PC2, principal component 2.

**Supplementary Material Table 6.** Data quality-related differences in the voxel-based morphometry in the secondary dataset (cluster-level FWE < 0.05).

| **Tissue class** | **MNI** | **Voxels** | **Location** | **Component** | **F-stat** |
| --- | --- | --- | --- | --- | --- |
| WM | -33, -2, -28 | 231 | L temporal lobe, sublobar | PC2 | 33.02 |
| WM | -15, 26, -2 | 111 | L frontal lobe, sublobar | PC2 | 25.00 |
| GM | -33, -1, -25 | 206 | L amygdala | PC2 | 34.38 |

Abbreviations: WM, white matter; GM, grey matter; L, left; PC2, principal component 2.

**Supplementary Material Table 7.** Results of the replication analysis of the clusters found significant on the whole-brain level in the primary cohort (cluster-level FWE < 0.05). Bolded false discovery rate (FDR) values indicate successful replication. Effect size in the form of Cohen’s d is provided only for the replicated findings.

| **Tissue class** | **MNI** | **Voxels** | **Location** | **Directionality** | **t-stat** | **FDR** | **Cohen’s d** |
| --- | --- | --- | --- | --- | --- | --- | --- |
| GM | -6, -47, 4 | 213 | B retrosplenial cortex | M > E | 3.80 | **0.004** | -0.633 |
| WM | 38, -30, 41 | 101 | R parietal lobe, sublobar | M > E | 3.41 | **0.006** | -0.568 |
| GM | 44, -42, -38 | 89 | R cerebellar lobule VIIa crus I  R inferior temporal gyrus | M > E | 1.44 | 0.305 |  |
| WM | 8, 3, -2 | 365 | R internal capsule | E > M | 1.26 | 0.305 |  |
| WM | -28, -72, 22 | 95 | L parietal lobe, sublobar | M > E | 1.26 | 0.305 |  |
| GM | 8, -11, -1 | 123 | R thalamus | M > E | 0.91 | 0.427 |  |
| GM | 2, -85, -34 | 151 | R cerebellar lobule VIIa crus II  R occipital pole | E > M | 0.55 | 0.584 |  |

Abbreviations: WM, white matter; GM, grey matter; B, bilateral; R, right; L, left; M, morning; E, evening.

**Supplementary Material Table 8.** Results of the replication analysis of the clusters found significant on the whole-brain level in the secondary cohort (cluster-level FWE < 0.05). Bolded false discovery rate (FDR) values signify successful replication, while italics indicate uncorrected p values < 0.05. Effect size in the form of Cohen’s d is provided only for the replicated findings.

| **Tissue class** | **MNI** | **Voxels** | **Location** | **Directionality** | **t-stat** | **FDR** | **Cohen’s d** |
| --- | --- | --- | --- | --- | --- | --- | --- |
| WM | 33, 0, -38 | 174 | R temporal, sublobar | M > E | 3.85 | **0.018** | -0.641 |
| GM | -33, -6, -15 | 143 | L amygdala  L hippocampus | E > M | 3.19 | **0.035** | 0.531 |
| WM | -31, -8, -15 | 226 | L temporal, sublobar | M > E | 3.18 | **0.035** | -0.530 |
| WM | -30, -29, 49 | 136 | L parietal, sublobar | M > E | 3.09 | **0.035** | -0.515 |
| WM | 26, -35, 45 | 630 | R parietal, sublobar | M > E | 2.97 | **0.038** | -0.495 |
| WM | 47, -4, 29 | 172 | R parietal, subgyral | M > E | 2.74 | *0.053* |  |
| WM | -18, 32, -16 | 123 | L orbitofrontal | M > E | 2.71 | *0.053* |  |
| WM | -21, -39, 55 | 124 | L parietal, sublobar | M > E | 2.56 | *0.065* |  |
| WM | -28, -44, 37 | 112 | L parietal, sublobar | M > E | 2.47 | *0.065* |  |
| WM | -22, -63, 43 | 134 | L parietal, subgyral | M > E | 2.46 | *0.065* |  |
| WM | 17, -5, 53 | 101 | R frontal, subgyral | M > E | 2.44 | *0.065* |  |
| WM | -19, 41, -7 | 97 | L orbitofrontal | M > E | 2.31 | *0.081* |  |
| GM | 8, 30, 19 | 119 | B anterior cingulate cortex | M > E | 1.91 | 0.177 |  |
| WM | 18, 7, 49 | 187 | R parietal, subgyral | M > E | 1.71 | 0.247 |  |
| WM | 11, -56, 66 | 127 | R parietal, subgyral | M > E | 1.64 | 0.262 |  |
| WM | 39, -50, 17 | 110 | R temporoparietal junction | M > E | 1.61 | 0.263 |  |
| GM | 15, -76, -71 | 310 | R cerebellar cortex | E > M | 1.49 | 0.307 |  |
| WM | -13, -5, 60 | 697 | L parietal, subgyral | M > E | 1.32 | 0.390 |  |
| WM | -37, -80, -22 | 152 | L occipital, subgyral | M > E | 1.25 | 0.415 |  |
| GM | -15, -83, -64 | 128 | L cerebellar cortex | E > M | 1.07 | 0.525 |  |
| WM | -16, 46, 26 | 134 | L frontal, sublobar | M > E | 1.04 | 0.525 |  |
| GM | -10, -89, 2 | 98 | B cuneus | E > M | 0.97 | 0.553 |  |
| WM | -30, -1, 36 | 96 | L frontal, subgyral | M > E | 0.89 | 0.596 |  |
| WM | -34, -61, -11 | 375 | L occipital, subgyral | M > E | 0.86 | 0.596 |  |
| WM | -10, -32, -19 | 143 | Brainstem | E > M | 0.57 | 0.819 |  |
| WM | 15, -11, 68 | 134 | R frontal, subgyral | M > E | 0.53 | 0.819 |  |
| GM | -34, -54, -21 | 116 | L fusiform gyrus | E > M | 0.50 | 0.819 |  |
| GM | 61, -6, -15 | 1003 | R middle temporal gyrus | E > M | 0.48 | 0.819 |  |
| GM | -39, -80, -57 | 98 | L cerebellar cortex | E > M | 0.43 | 0.828 |  |
| GM | 58, -31, -5 | 105 | R middle temporal gyrus | E > M | 0.40 | 0.828 |  |
| GM | 42, -59, 27 | 103 | R angular gyrus | E > M | 0.25 | 0.902 |  |
| GM | -62, -24, -9 | 572 | L middle temporal gyrus | E > M | 0.25 | 0.902 |  |
| GM | -30, -64, -11 | 118 | L fusiform gyrus | E > M | 0.20 | 0.906 |  |
| GM | 0, 6, 70 | 105 | B superior frontal gyrus | M > E | 0.18 | 0.906 |  |
| GM | 0, 63, 0 | 720 | B frontal pole  L orbitofrontal cortex | M > E | 0.09 | 0.953 |  |
| GM | -37, -80, -22 | 215 | L inferior occipital gyrus | E > M | 0.04 | 0.972 |  |

Abbreviations: WM, white matter; GM, grey matter; B, bilateral; R, right; L, left; M, morning; E, evening.

**Supplementary Material Figure 1.** The distribution of signal-to-noise ratio (SNR) across the dataset.

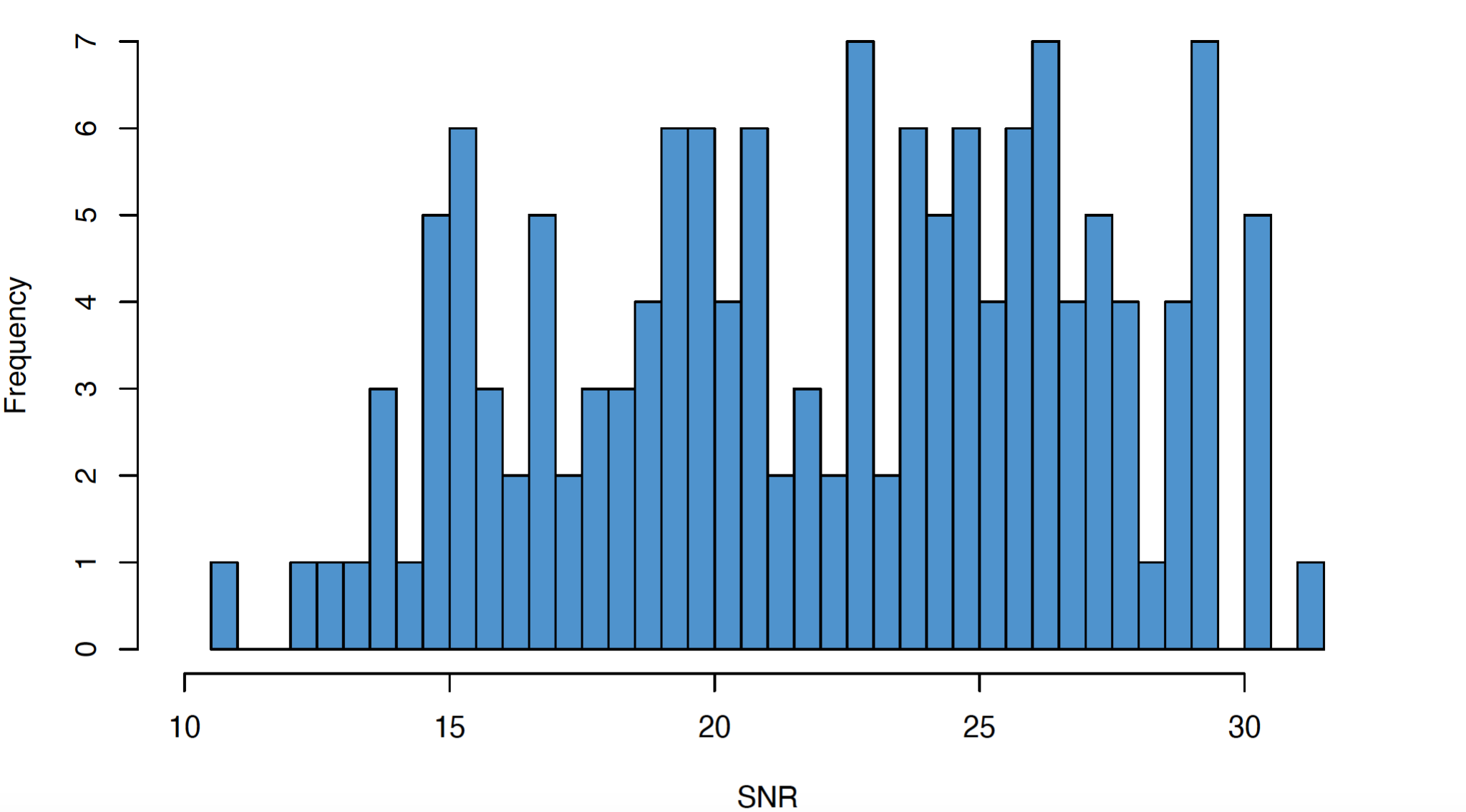

**Supplementary Material Figure 2.** The distribution of contrast-to-noise (CNR) across the dataset.

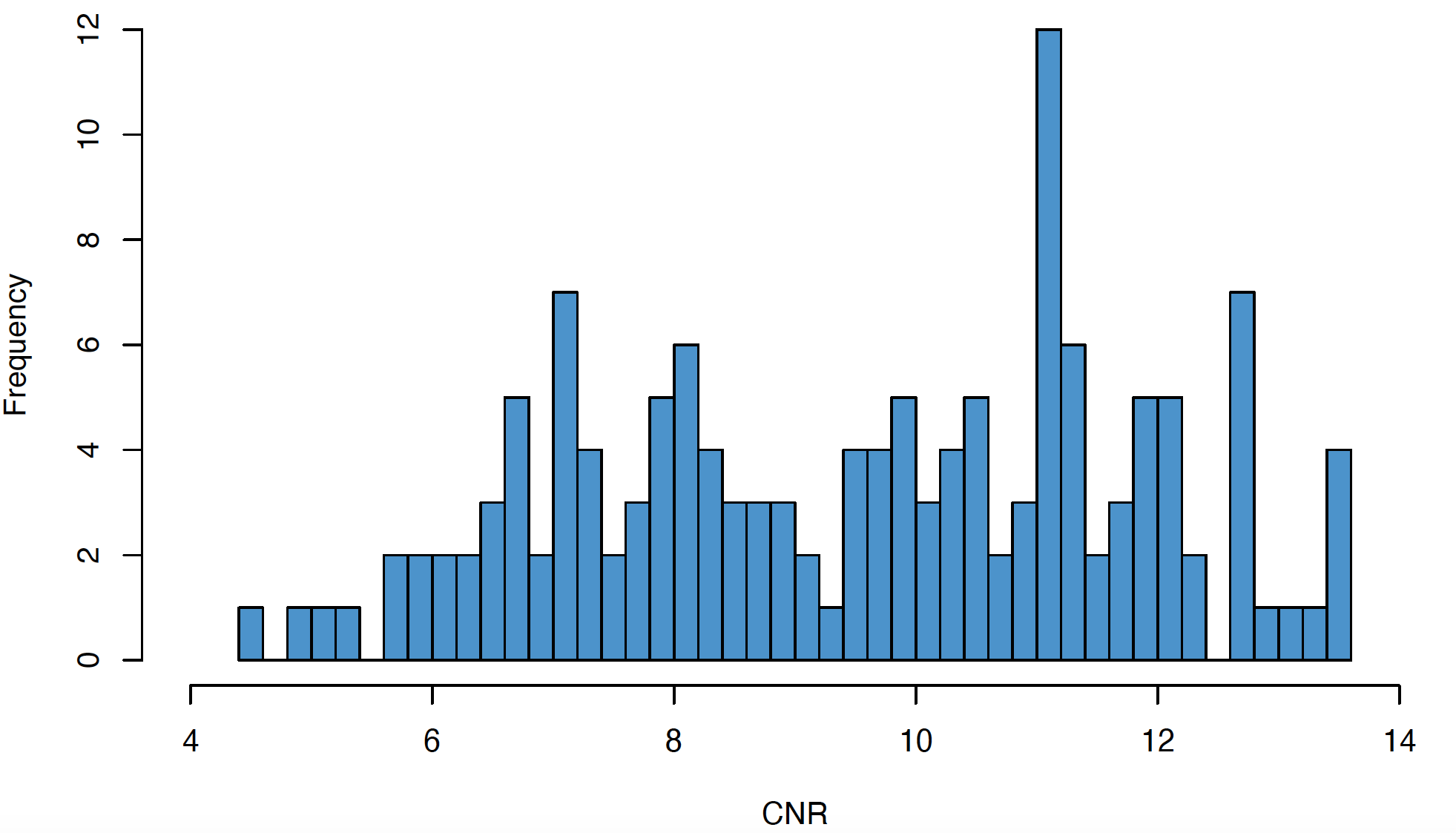

**Supplementary Material Figure 3.** The distribution of foreground-to-background energy ratio (FBER) across the dataset.

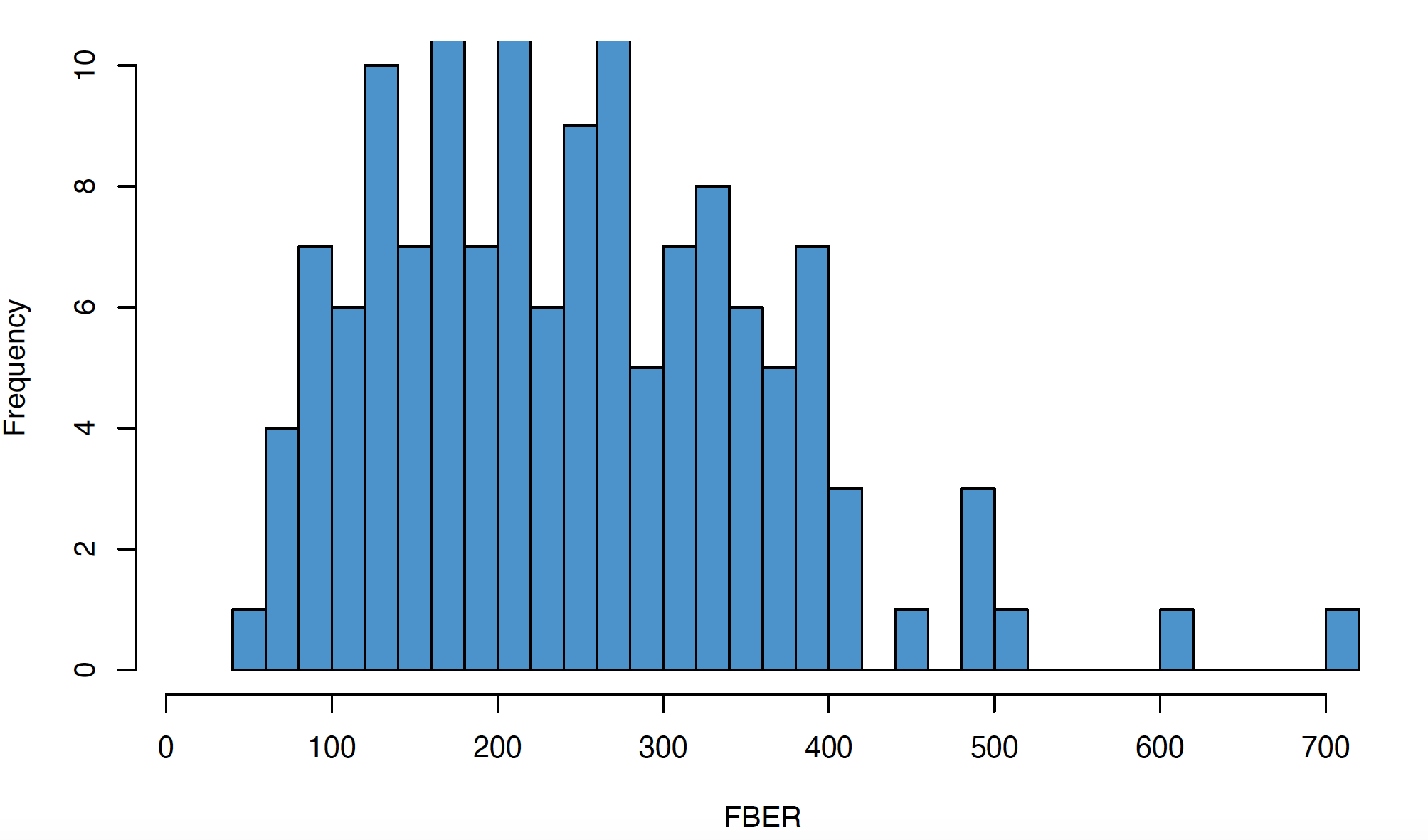

**Supplementary Material Figure 4.** The distribution of voxel smoothness defined as full width half maximum (FWHM) across the dataset.

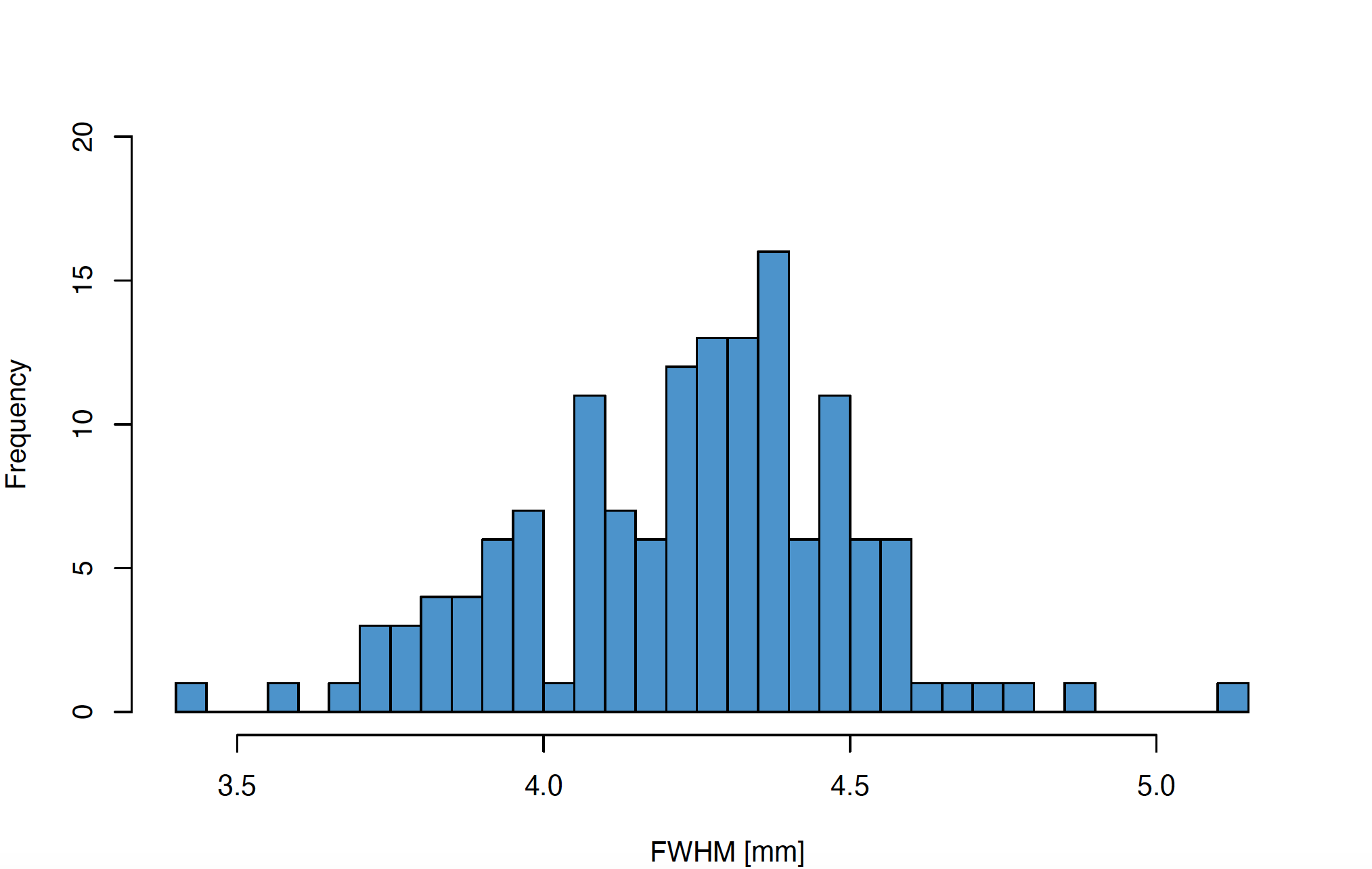

**Supplementary Material Figure 5.** The distribution of entropy focus criterion (EFC) across the dataset.

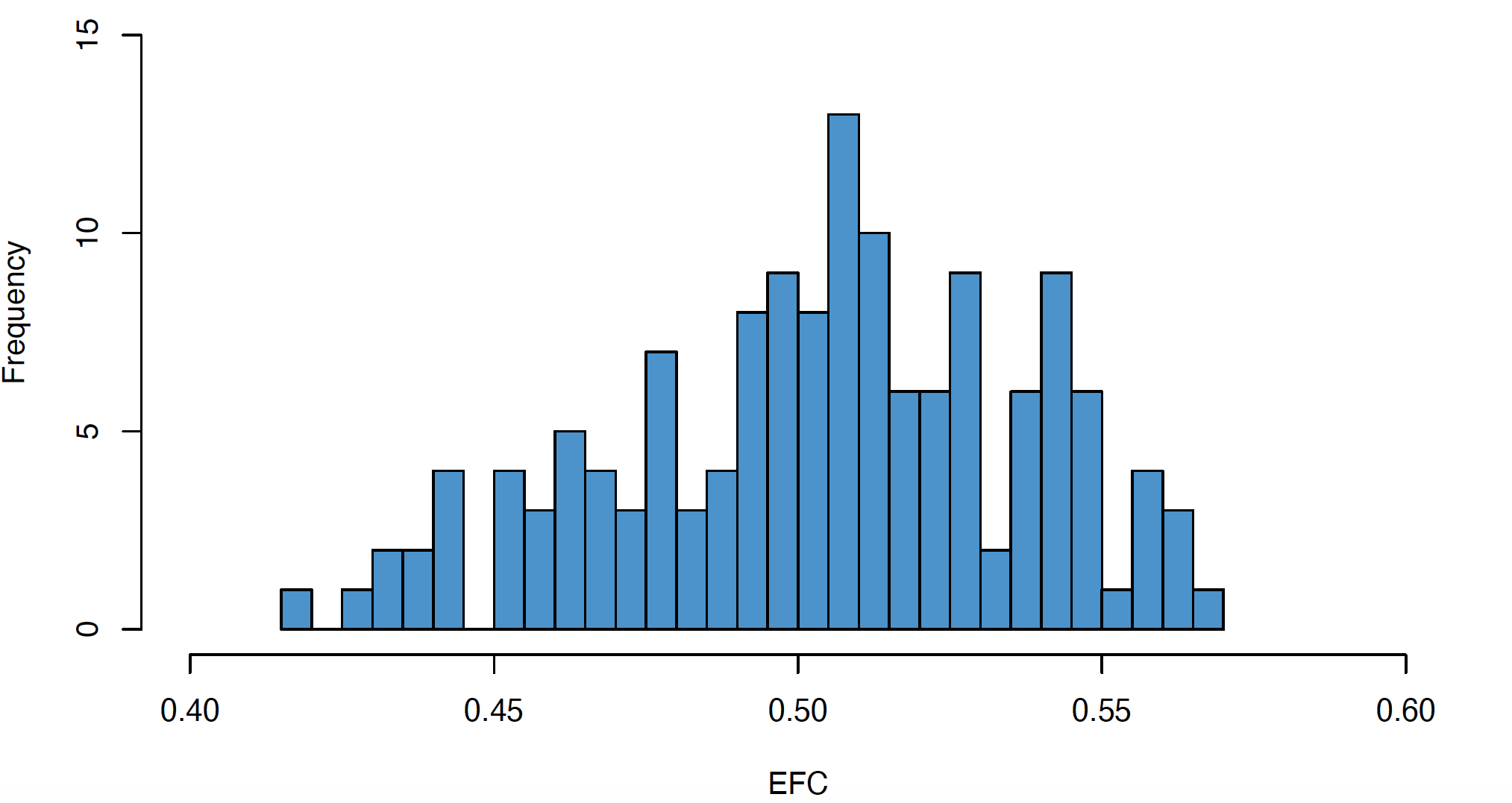

**Supplementary Material Figure 6.** The distribution of fraction of artefact background voxels (QI1) across the dataset.

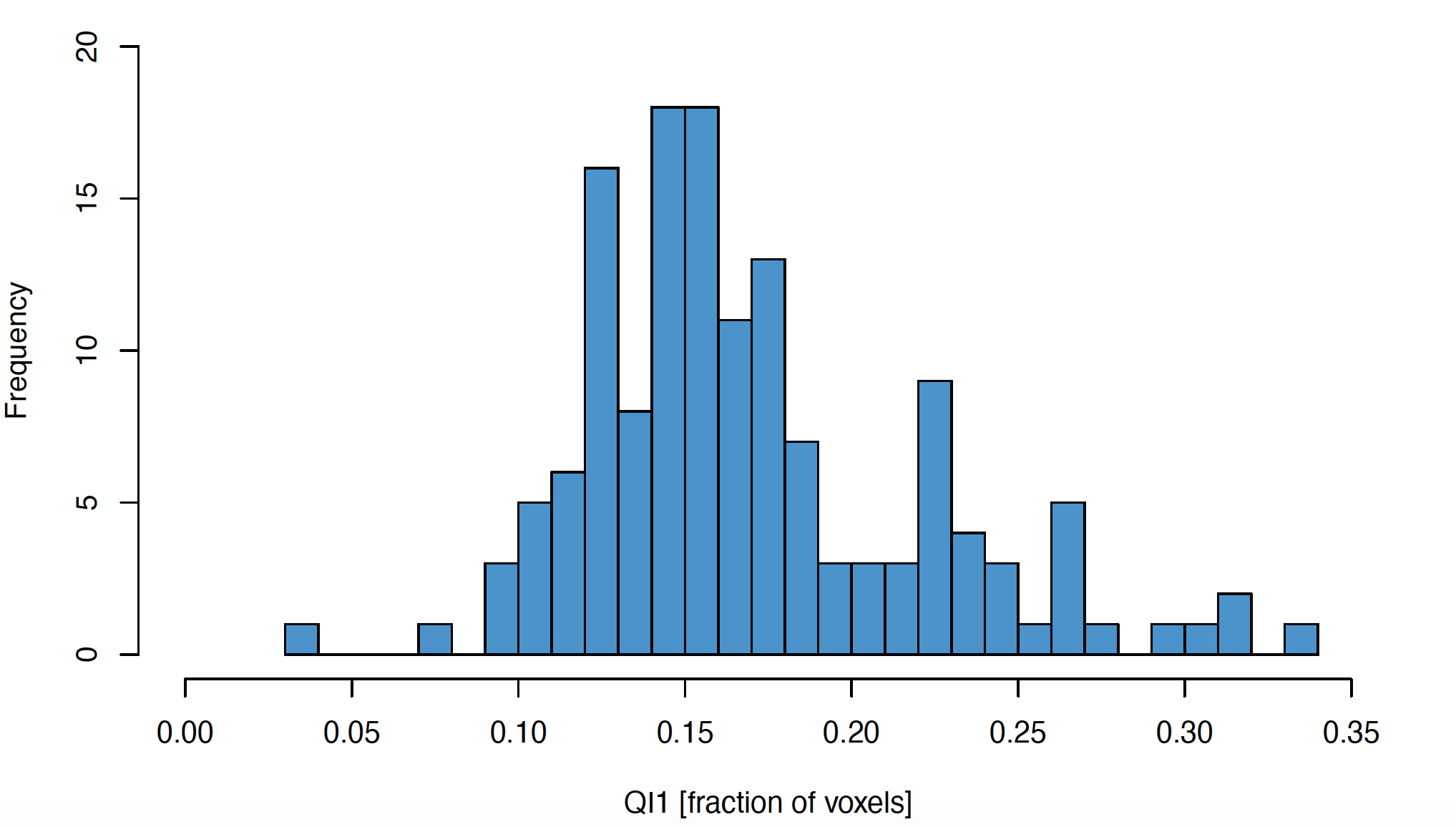

**Supplementary Material Figure 7.** The variability in VBM associated with data quality in the primary dataset (cluster-level FWE < 0.05). Principal component 1 (PC1) was related to signal-to-noise ratio, contrast-to-noise ratio and foreground-to-background energy ratio. Principal component 2 (PC2) was associated with entropy focus criterion, a measure related to motion and ghosting artefacts, and fraction of artefact background voxels. Abbreviations: GM, grey matter; WM, white matter.

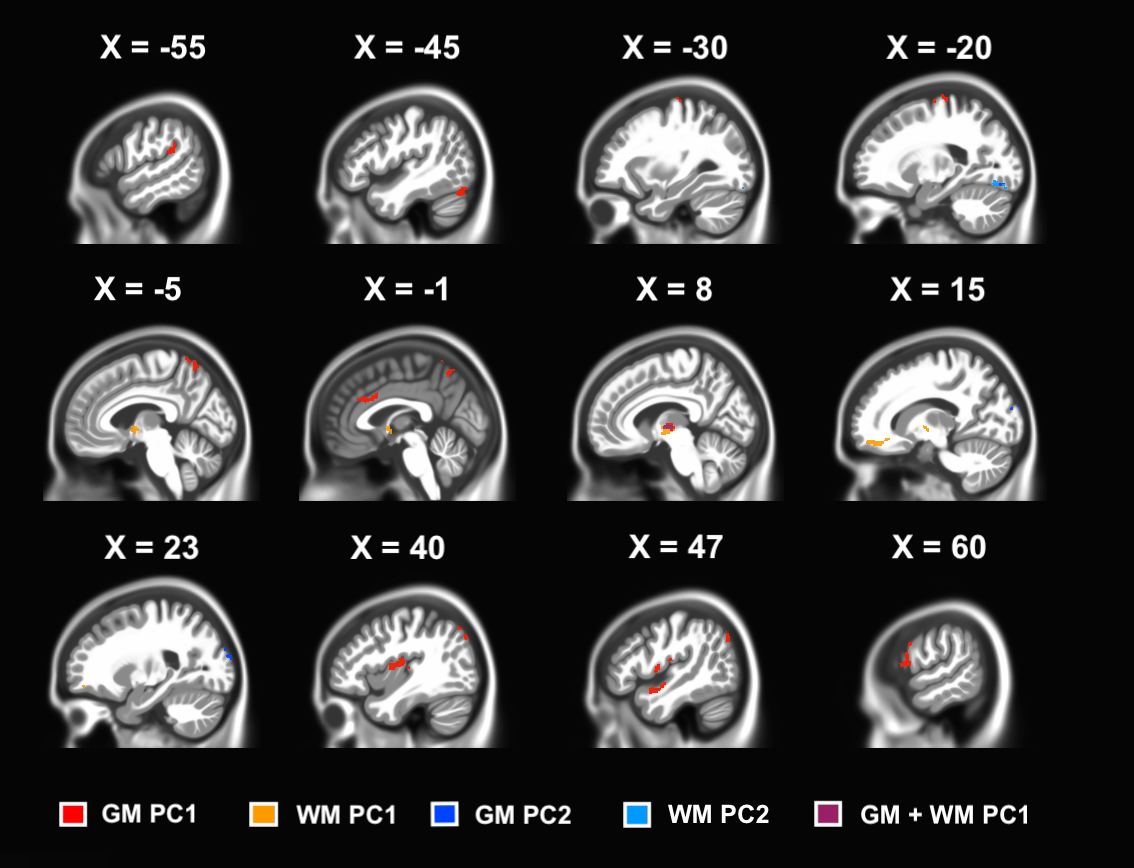

**Supplementary Material Figure 8.** The variability in voxel-based morphometry (VBM) associated with data quality in the secondary dataset (cluster-level FWE < 0.05). Principal component 2 (PC2) was associated with entropy focus criterion, a measure related to motion and ghosting artefacts, and fraction of artefact background voxels. Abbreviations: GM, grey matter; WM, white matter.

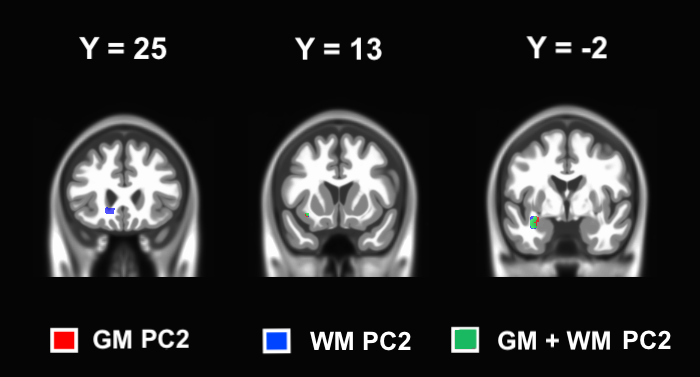

**Supplementary Material Figure 9.** The spatial overlap between the morning-to-evening decreases in white matter (WM) volume in the right parietal lobe found in the two analyses. The name of the dataset implies which whole-brain analysis the mask was taken from.

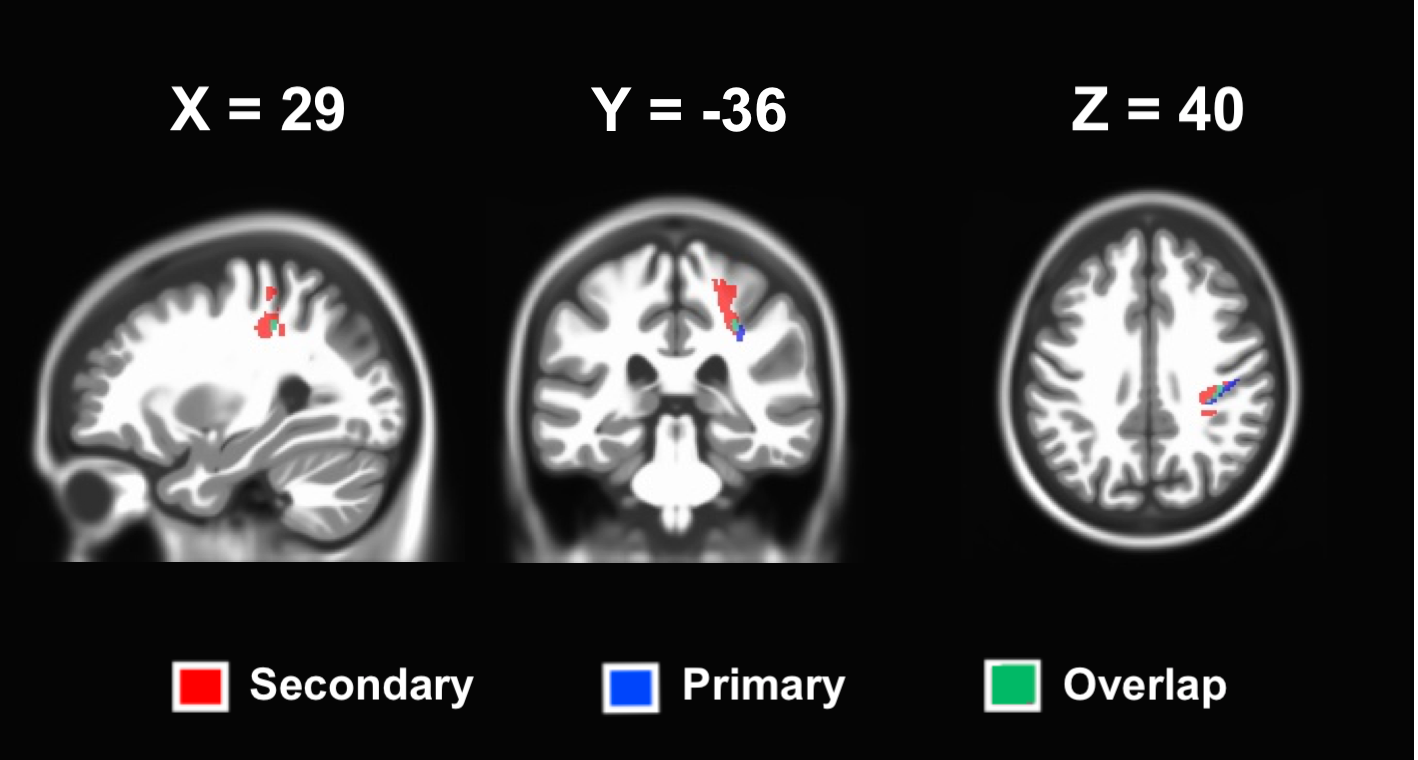
